# MorphQ: label-free quantification and visualisation of complex morphology from standardised specimen images

**DOI:** 10.64898/2026.08.11.744091

**Authors:** Ying-Yu Chen, Guan-Shuo Mai, Dustin R. Rubenstein, Chia-Hsuan Wei, Sheng-Feng Shen

## Abstract

1. Quantifying complex morphology from images remains difficult because predefined descriptors capture only selected traits. Yet, supervised machine learning models for images require labels and often produce task-specific features that are hard to interpret as biological traits.
2. We present MorphQ, a label-free, self-supervised method that learns a quantitative morphospace from standardised specimen images. Its encoder produces feature vectors for statistical analysis, and its decoder converts analysed positions in morphospace into human-interpretable images, including hypothetical forms not represented by sampled specimens or sampled taxa.
3. Using 1,868 Lepidoptera species, we tested whether MorphQ’s label-free features were more useful for downstream analysis than features from principal component analysis (PCA) or a supervised species-classification machine learning model. As a diagnostic probe of downstream biological utility, MorphQ features supported higher low-label family-classification accuracy than comparator features, and retained stronger family-level similarity for species absent from model training, indicating better generalisation to species not seen during model training.
4. Two case studies link MorphQ morphospaces to species-level elevation and assemblage-level functional diversity while keeping statistical patterns visually inspectable. MorphQ provides a reproducible framework for constructing interpretable morphological trait spaces when predefined descriptors are incomplete and labelled data are limited.

Data/code for peer review: An anonymised repository containing the source code, trained model weights, example data, configuration files and scripts required to reproduce the analyses is available at https://anonymous.4open.science/r/MorphQ-ECD4/.

## Introduction

Organismal morphology provides the basis for many ecological and evolutionary questions, yet objectively quantifying complex form remains a persistent methodological challenge. Researchers need comparable morphological features to study how sexual selection shapes traits, how organisms vary across geographic and environmental gradients, and how morphological disparity differs among lineages and through evolutionary time (Wallace, 1878; Roy & Foote, 1997; Klingenberg, 2010; Swenson, 2012; Moczek & Emlen, 2000; Tian et al., 2016; Briggs et al., 1992; Wills et al., 1994). Each of these questions requires a morphospace in which organisms can be compared quantitatively. Gould (1991) emphasised that the challenge lies not only in measuring particular traits, but also in defining a morphospace that can describe different organisms in a comparable and interpretable way.

The typical solution has been to define measurable traits before analysis, a strategy that has produced powerful tools in geometric morphometrics, colouration analysis, and pattern quantification but also imposes an important constraint: the final trait space is determined a priori by the specific features that researchers choose in advance (Cooke & Terhune, 2015; Cervantes et al., 2016; White et al., 2015; Kemp et al., 2015). Colour analyses often begin by selecting particular body regions and summarising reflectance or pixel statistics within them, whereas shape and pattern analyses typically require predefined landmarks, outlines, areas, or indices (Stevens et al., 2007; Dale et al., 2015; Pérez-Rodríguez et al., 2017; Chan et al., 2019; Pike, 2018; Stoddard & Osorio, 2019). These approaches can be reproducible once the descriptors have been defined, but they generally capture only selected aspects of the image. They may miss co-varying combinations of colour, shape, size, stripes, spots, and other visual information that are obvious to human observers but difficult to specify as separate measurements.

This limitation resembles the feature-engineering problem that long constrained machine learning. Earlier algorithms often depended on human-designed descriptors (Ojala et al., 1996; Lowe, 1999; Dalal & Triggs, 2005), until deep learning allowed models to learn image representations directly from data (Hinton et al., 2006; Hinton & Salakhutdinov, 2006; Krizhevsky et al., 2012; LeCun et al., 2015; Goodfellow et al., 2016). For ecological and evolutionary biology, this development is important because images contain abundant unstructured information that may be relevant to biological traits but that are difficult to quantify manually (Wäldchen & Mäder, 2018). In principle, a model that derives visual representations directly from image variation could help construct a morphospace that depends less on a priori decisions about which trait components should be measured.

Despite the obvious potential of machine learning approaches to studies of organismal morphology, supervised deep learning does not fully solve the problem of morphological trait quantification. Supervised models require many labelled examples to cover the data distribution, a requirement that is often unrealistic in ecological studies, where many taxa often have few standardised images or sparse annotations (LeCun et al., 2015; Goodfellow et al., 2016). The deeper limitation is that the representations learned by supervised models are guided by the recognition task (Zhong et al., 2016). A species classifier, for example, may learn features similar to a dichotomous key: useful for distinguishing labels, but not necessarily providing comprehensive descriptions of morphology. A model may also exploit biases in backgrounds, lighting, or image acquisition when those cues help classification, even if they are unrelated to the biological traits of interest. As a result, high classification accuracy does not, by itself, indicate that the resulting feature vectors form a general, interpretable, and reusable feature space, let alone a biologically meaningful morphospace.

Image-based morphological analysis for ecological and evolutionary research therefore needs to meet a different set of requirements. It should quantify morphology without requiring extensive labels or manually predefined descriptors; reduce high-dimensional images to a feature space that can be used in statistical analyses; preserve biologically meaningful similarity among organisms; and provide a means of inspecting what the quantitative features represent. This matters because the same feature space may be reused in different downstream analyses, such as comparisons among taxa, tests of trait-environment relationships, and analyses of functional diversity.

Here, we present MorphQ, a label-free, self-supervised representation-learning framework designed to turn image collections into quantitative and visually interpretable morphospaces (see Box 1 for a glossary of machine learning terms). It is intended for standardised, consistently aligned specimen images in which the focal organismal structure is positioned comparably across images. MorphQ is particularly suited to datasets with many taxa or groups of organisms, but few standardised images per taxon, where supervised training within each category is impractical and when manually predefined descriptors are likely to capture only part of the relevant morphology.

### Box 1. Glossary of machine-learning terms

Terms are defined at their first use in the main text, but collected here for reference. Definitions describe each term operationally, relating it to familiar methods in ecology and evolution where possible.

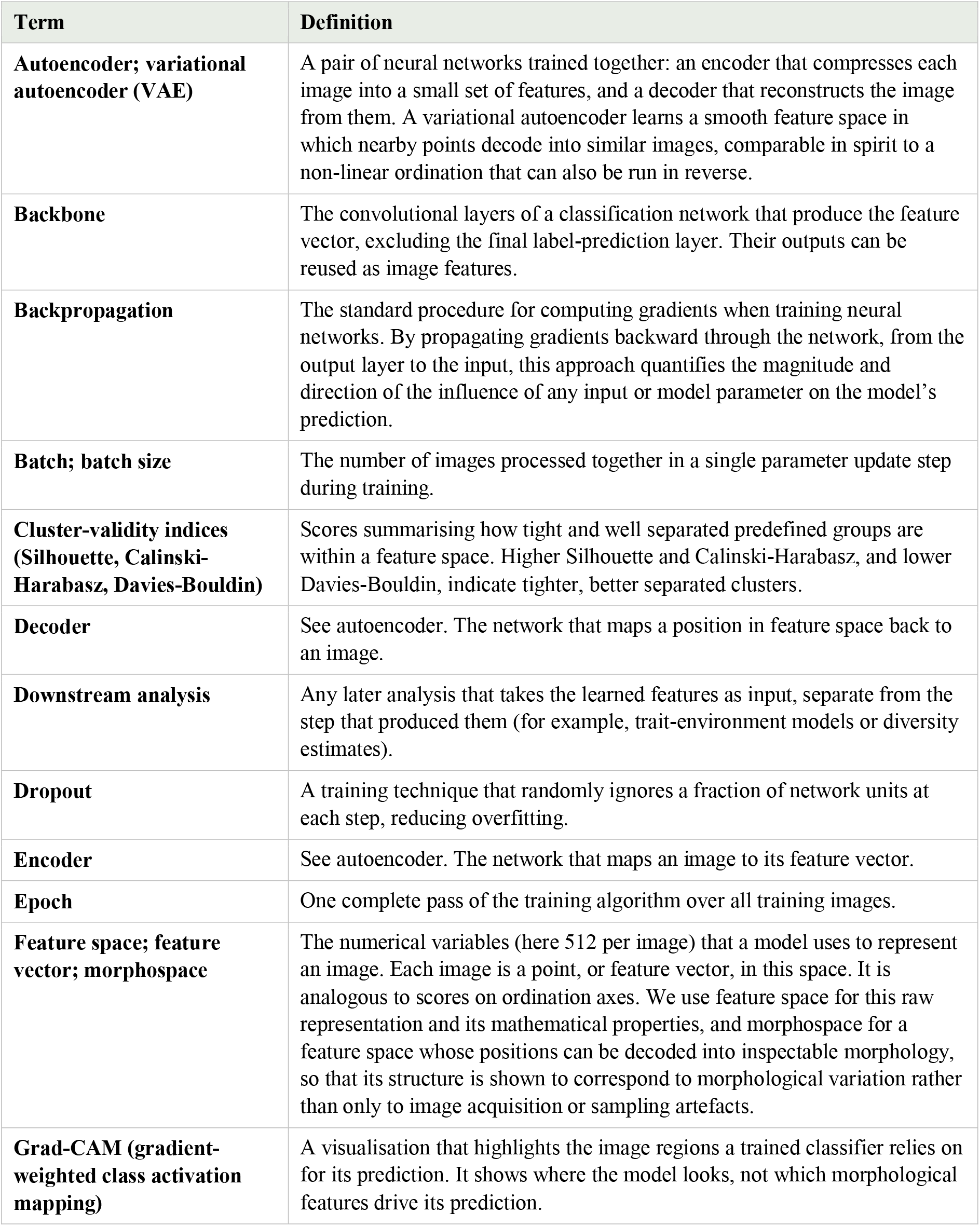
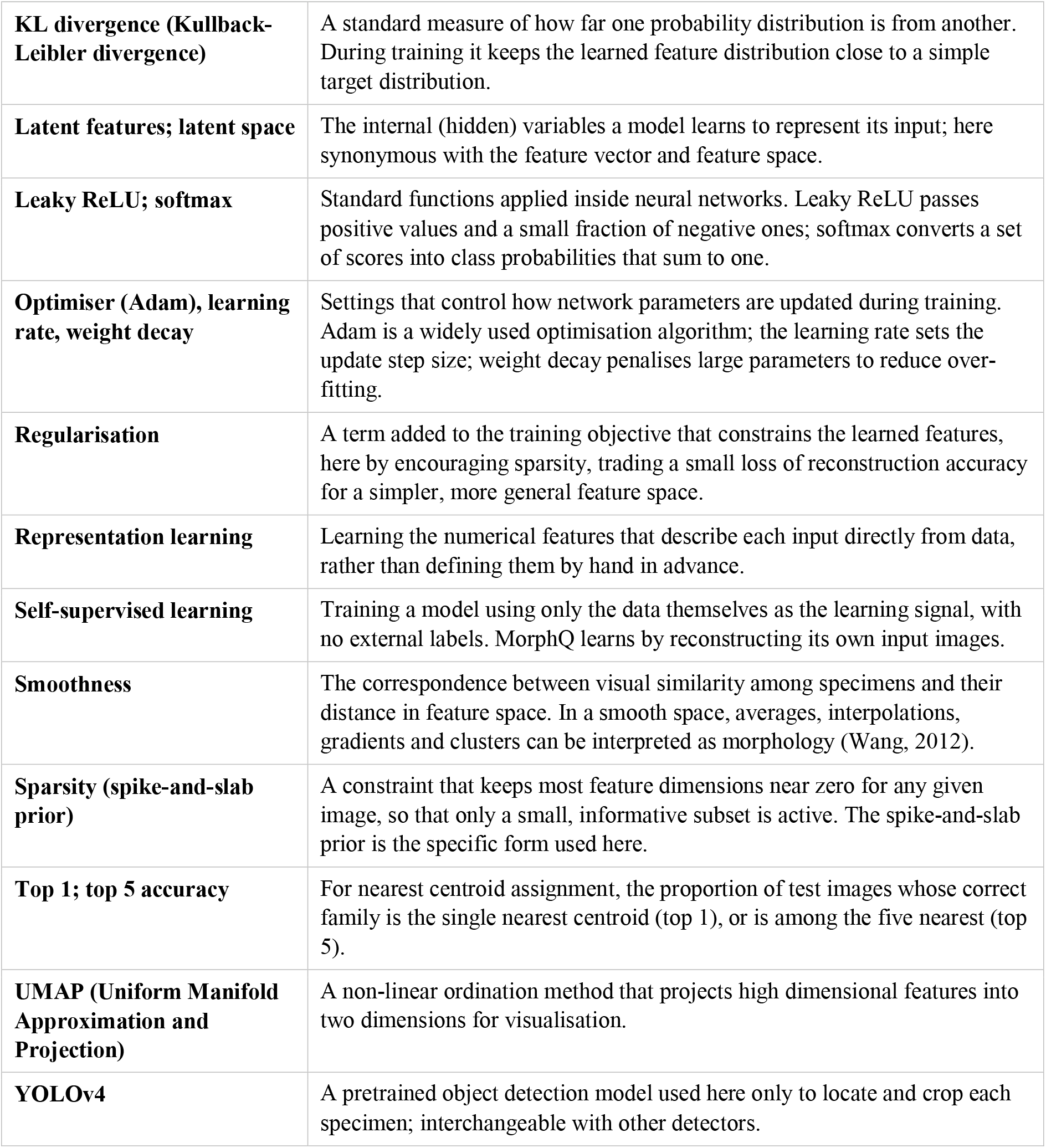

| Term | Definition |
| --- | --- |
| **Autoencoder; variational autoencoder (VAE)** | A pair of neural networks trained together: an encoder that compresses each image into a small set of features, and a decoder that reconstructs the image from them. A variational autoencoder learns a smooth feature space in which nearby points decode into similar images, comparable in spirit to a non-linear ordination that can also be run in reverse. |
| **Backbone** | The convolutional layers of a classification network that produce the feature vector, excluding the final label-prediction layer. Their outputs can be reused as image features. |
| **Backpropagation** | The standard procedure for computing gradients when training neural networks. By propagating gradients backward through the network, from the output layer to the input, this approach quantifies the magnitude and direction of the influence of any input or model parameter on the model's prediction. |
| **Batch; batch size** | The number of images processed together in a single parameter update step during training. |
| **Cluster-validity indices (Silhouette, Calinski-Harabasz, Davies-Bouldin)** | Scores summarising how tight and well separated predefined groups are within a feature space. Higher Silhouette and Calinski-Harabasz, and lower Davies-Bouldin, indicate tighter, better separated clusters. |
| **Decoder** | See autoencoder. The network that maps a position in feature space back to an image. |
| **Downstream analysis** | Any later analysis that takes the learned features as input, separate from the step that produced them (for example, trait-environment models or diversity estimates). |
| **Dropout** | A training technique that randomly ignores a fraction of network units at each step, reducing overfitting. |
| **Encoder** | See autoencoder. The network that maps an image to its feature vector. |
| **Epoch** | One complete pass of the training algorithm over all training images. |
| **Feature space; feature vector; morphospace** | The numerical variables (here 512 per image) that a model uses to represent an image. Each image is a point, or feature vector, in this space. It is analogous to scores on ordination axes. We use feature space for this raw representation and its mathematical properties, and morphospace for a feature space whose positions can be decoded into inspectable morphology, so that its structure is shown to correspond to morphological variation rather than only to image acquisition or sampling artefacts. |
| **Grad-CAM (gradient-weighted class activation mapping)** | A visualisation that highlights the image regions a trained classifier relies on for its prediction. It shows where the model looks, not which morphological features drive its prediction. |
| **KL divergence (Kullback-Leibler divergence)** | A standard measure of how far one probability distribution is from another. During training it keeps the learned feature distribution close to a simple target distribution. |
| **Latent features; latent space** | The internal (hidden) variables a model learns to represent its input; here synonymous with the feature vector and feature space. |
| **Leaky ReLU; softmax** | Standard functions applied inside neural networks. Leaky ReLU passes positive values and a small fraction of negative ones; softmax converts a set of scores into class probabilities that sum to one. |
| **Optimiser (Adam), learning rate, weight decay** | Settings that control how network parameters are updated during training. Adam is a widely used optimisation algorithm; the learning rate sets the update step size; weight decay penalises large parameters to reduce over-fitting. |
| **Regularisation** | A term added to the training objective that constrains the learned features, here by encouraging sparsity, trading a small loss of reconstruction accuracy for a simpler, more general feature space. |
| **Representation learning** | Learning the numerical features that describe each input directly from data, rather than defining them by hand in advance. |
| **Self-supervised learning** | Training a model using only the data themselves as the learning signal, with no external labels. MorphQ learns by reconstructing its own input images. |
| **Smoothness** | The correspondence between visual similarity among specimens and their distance in feature space. In a smooth space, averages, interpolations, gradients and clusters can be interpreted as morphology (Wang, 2012). |
| **Sparsity (spike-and-slab prior)** | A constraint that keeps most feature dimensions near zero for any given image, so that only a small, informative subset is active. The spike-and-slab prior is the specific form used here. |
| **Top 1; top 5 accuracy** | For nearest centroid assignment, the proportion of test images whose correct family is the single nearest centroid (top 1), or is among the five nearest (top 5). |
| **UMAP (Uniform Manifold Approximation and Projection)** | A non-linear ordination method that projects high dimensional features into two dimensions for visualisation. |
| **YOLOv4** | A pretrained object detection model used here only to locate and crop each specimen; interchangeable with other detectors. |

To assess whether MorphQ met the requirements outlined above, we operationalised them as four criteria for downstream ecological and evolutionary analyses. We evaluated MorphQ-constructed feature spaces by comparing them with feature spaces produced by principal component analysis (PCA) and a supervised species-classification machine learning model. Specifically, we asked whether the resulting features could be reused in downstream tasks whose targets differed from their upstream fitting or training purpose; whether they formed a compact and less redundant feature space; whether they remained applicable beyond the fitted data; and whether their quantitative structure could be connected back to visual morphology. Finally, two case studies, at the species and assemblage levels, illustrate how MorphQ takes an analysis from images through to quantitative traits and back to inspectable morphology.

## Materials and methods

### Overview of MorphQ

MorphQ comprises two parts, an encoder and a decoder (Figure 1). The encoder quantifies morphological information in images into feature vectors, which supply numerical variables for downstream statistical analyses. The decoder can reconstruct or generate images from any feature combination, allowing the results of analyses, such as group summaries, interpolations, or trends along environmental gradients, to be rendered as images that humans can inspect and interpret biologically. MorphQ training requires only images. Metadata such as taxonomic, ecological, or environmental information are optional for validation or downstream analyses and are not involved in the learning process.

**Figure 1.**
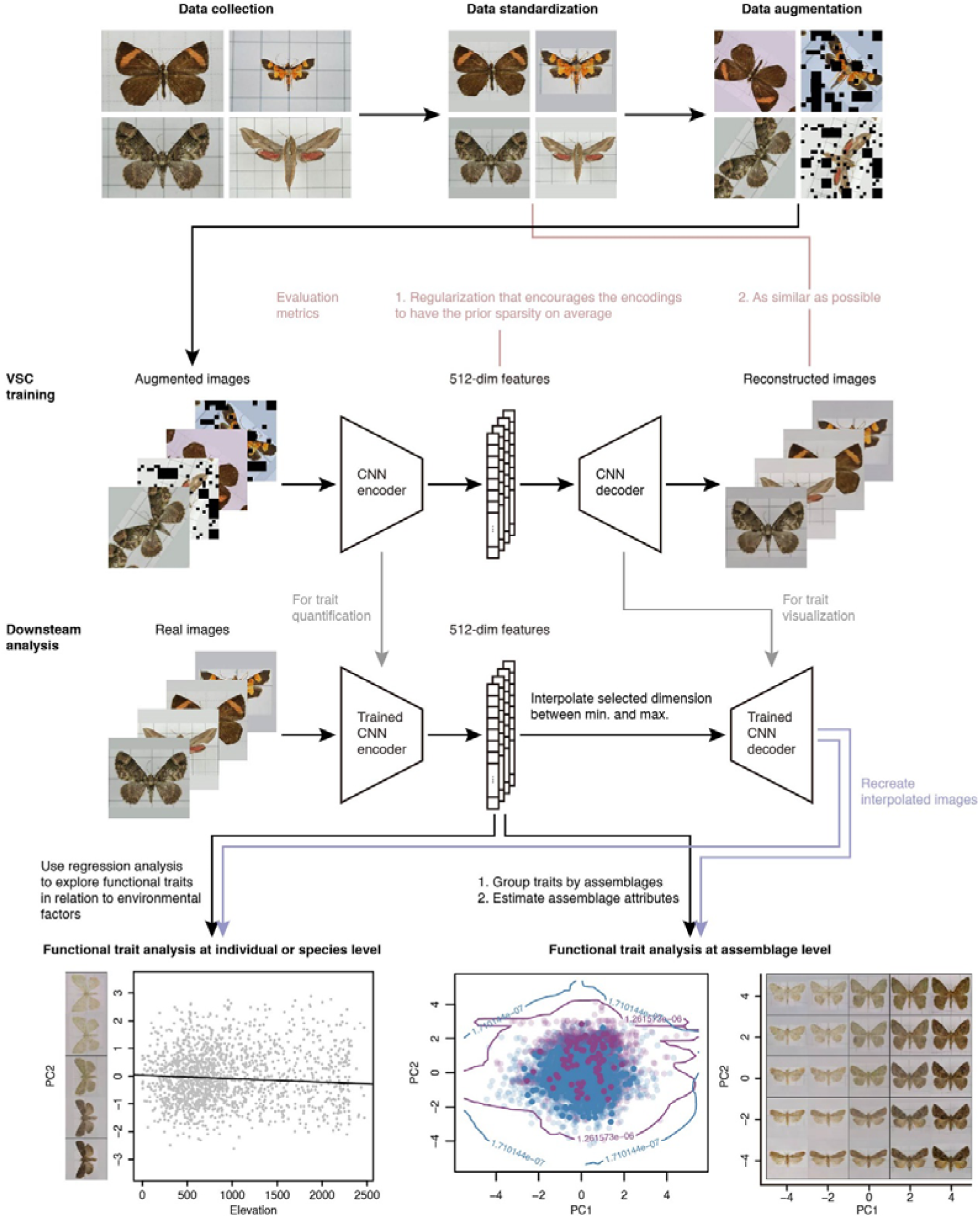
Overview for MorphQ. Steps comprise image standardisation, model training or application, feature extraction, optional aggregation by species or group, downstream analysis, and visualisation of specified morphospace positions as images.

Throughout, we distinguish the feature space from the morphospace. We use feature space for the raw 512-dimensional representation produced by an extractor and its mathematical properties (e.g., compactness, redundancy, and distances between points). We reserve morphospace for a feature space whose positions can be decoded into inspectable morphology, so that its structure is shown to correspond to morphological variation rather than only to image acquisition or sampling artefacts. Under this usage, any extractor yields a feature space, but a feature space becomes a morphospace only once this visual correspondence is established. To evaluate MorphQ-constructed feature spaces, we built three sets of 512-dimensional features from the same images using the MorphQ encoder and two comparator models, and assessed them against four criteria defined below.

### Image datasets and preprocessing

To evaluate MorphQ under both the intended standardised image setting and a small-sample scenario, and to test whether the resulting feature space remained applicable beyond the fitted data, we used three Lepidoptera specimen image datasets with corresponding metadata, each serving a different purpose. First, the primary dataset contained 3,736 images from 1,868 species (two images per species), and was used to train MorphQ and conduct the main feature evaluation analyses. Second, the reference dataset was a separate large dataset containing 68,913 labelled images and was used only to train the large-sample DeepClassifier reference for the downstream task comparison; it was neither used to train MorphQ nor to generate the main MorphQ-derived features. Third, the isolated dataset contained 385 species (one image per species), and was kept completely unseen during all training and fitting processes to test the third criterion.

The metadata of the primary dataset were based on the digitised moth specimens curated by the Taiwan Moth Information Center (TESRI) on the open access GBIF “Dataset of Moth Specimen from TESRI” (https://doi.org/10.15468/kjjlnf, accessed in August 2019). For each species, two specimen images were retained.

For the large labelled reference dataset, we assembled additional specimen images downloaded from the official TESRI site (https://twmoth.tesri.gov.tw, accessed in February 2022), targeting only the species represented in the primary dataset. After quality filtering (see Appendix S1 in Supporting Information), 68,913 Lepidoptera specimen images were retained.

The isolated dataset was also downloaded from TESRI and contained 385 species not represented in the primary dataset, with one image per species. Details of this comparison are given in Appendix S1.

Images in all datasets were standardised using the same cropping, resizing, and padding workflow. Specimens were detected and cropped automatically with a YOLOv4 model (Bochkovskiy et al. 2020), and each cropped image was rescaled and padded to a 256 × 256 × 3 RGB image with a background-matched padding colour (Appendix S1).

### MorphQ model design and training objective

Using the primary dataset, we trained MorphQ to learn a feature space from standardised specimen images. MorphQ was implemented following variational sparse-coding (VSC) and beta variational autoencoder (beta-VAE) principles (Kingma & Welling, 2014; Higgins et al., 2017; Tonolini et al., 2020). During training, the encoder received augmented images with introduced noise, while the decoder was trained to reconstruct the corresponding target images, that is, the augmented images without introduced noise. This denoising objective prevented the model from directly copying input pixels and instead required it to infer missing visual structure from the remaining image information (Vincent et al., 2010).

The encoder maps an input image into a 512-dimensional spike-and-slab latent feature distribution, described by three sets of 512-dimensional parameters: mean, log variance, and log spike, from which a 512-dimensional feature vector is sampled by reparametrisation. During training, the sampled 512-dimensional feature vector was fed to the decoder to reconstruct the corresponding denoised image.

Training minimised two loss components: the reconstruction loss and the regularisation term, with (L_total_ = L_rec_ + βL_reg_) and (β = 1). Reconstruction loss was calculated as the pixel-level sum of squared errors of RGB values between reconstructed images and the corresponding denoised images, averaged across the batch. The regularisation term was the spike-and-slab KL divergence between the learned feature distribution and a target spike-and-slab distribution centred on zero. The sparse parameter alpha was set to 0.01. This prior encouraged many feature dimensions to remain near zero while allowing a subset of dimensions to vary among images, producing a sparse feature space.

The encoder followed a convolutional down-sampling architecture, in which a stack of residual blocks progressively expanded the channel width while compressing image information into the 512-dimensional latent feature distribution (Table S2). The decoder was mostly symmetric to the encoder, with down-sampling layers replaced by up-sampling layers (Table S2).

The augmentation pipeline comprised geometric and colour transformations, and noise introduction comprised random rotation and coarse dropout (Table S2). These augmentation and noise introduction steps were used only during training, to encourage the encoder to learn robust visual representations rather than to memorise individual images.

MorphQ was implemented in PyTorch and trained on the primary dataset using Adam optimisation (Table S2). No formal early stopping rule or predefined epoch limit was used; the final model was taken once both the reconstruction loss and the regularisation term had stabilised in the training record.

### Feature extraction and visualisation

After training, each image was passed through the trained MorphQ encoder to obtain a stochastic rather than deterministic feature vector. We therefore resampled 1,000 times from the parameters to obtain 1,000 feature vectors of 512 dimensions per image and averaged them dimension-wise to obtain a Monte Carlo estimate of the image-level feature vector. These averaged 512-dimensional vectors were used as MorphQ-derived morphological features in downstream analyses.

The decoder can map any given feature space position or feature combination, such as a group centroid or statistical gradient, back into a human-inspectable image that visualises that position rather than necessarily corresponding to a sampled specimen image. Outputs from downstream analyses, such as coordinates along principal-component axes in reduced feature spaces derived from the 512-dimensional MorphQ feature space, were first inverse-transformed back into the original 512-dimensional space and then decoded. Additional decoder-based visualisations included family centroids calculated from mean feature vectors and the key feature combinations that the family classifier relied on to predict families, selected as described below (Figure S3).

### Comparator methods

To interpret MorphQ-derived features under comparable output dimensionality, we used two main image-derived feature baselines: PCA and a small-sample DeepClassifier. PCA provided an unsupervised linear baseline based on pixel-level variation, whereas the small-sample DeepClassifier provided a supervised feature baseline trained for species recognition using the same small-sample primary dataset as MorphQ. These comparators were not intended to cover all possible morphological descriptors. Instead, they isolated two contrasts relevant to MorphQ: how a label-free autoencoder-derived feature space differs from a linear image embedding, and how it differs from a neural network feature representation optimised for a supervised recognition task.

PCA features were obtained by retaining 512 principal components of the flattened standardised images, matching the feature dimensionality used by MorphQ (Appendix S1); these 512 components explained 87.6% of the total variance. PCA was also the only decoder-like comparator, because PCA inverse transformation can project retained component values back into image space (Appendix S1). The large-sample DeepClassifier reference, trained on the reference dataset of 68,913 images, was applied to the primary dataset images only to evaluate how much the additional labelled images improved the supervised feature extractor; it was not used to train MorphQ, to fit PCA, or as a general comparator for the other criteria.

The small-sample DeepClassifier followed the MorphQ encoder architecture as closely as possible to reduce architectural differences between the supervised comparator and MorphQ. However, the variational latent head and decoder used by MorphQ were replaced by adaptive average pooling and a fully connected species classification head, so that the pooled representations formed 512-dimensional feature vectors directly comparable with MorphQ features (Appendix S1). Training details for the small-sample DeepClassifier, including the primary dataset training, validation and testing split ratio, optimisation settings and early stopping rule, are given in Appendix S1, together with those for the large-sample DeepClassifier reference.

### Evaluation of image-derived morphological representations

We evaluated MorphQ according to four criteria derived from representation learning, which we interpreted as requirements for an image-derived feature space in ecological and evolutionary analyses (Bengio et al., 2013; Tschannen et al., 2018). A useful feature space should be reusable in downstream analyses, compact and non-redundant, applicable beyond the fitted data, and translatable back into visual morphospace.

For the first criterion, downstream reusability, we used family classification as a diagnostic downstream probe of feature utility, not as the intended end use of MorphQ, asking whether a simple classifier could recover a higher level biological grouping from features learned under different upstream objectives. MorphQ, PCA and the small-sample DeepClassifier were therefore treated only as feature extractors: each generated 512-dimensional feature vectors for all 3,736 images in the primary dataset, and the same downstream family classifier architecture was trained separately on each feature set. This design separated feature learning from downstream evaluation, since MorphQ learned features without taxonomic labels, PCA provided a linear image embedding and the small-sample DeepClassifier learned features through supervised species prediction. The large-sample DeepClassifier reference was included only in this downstream probe, to evaluate whether additional labelled images improved the supervised feature extractor. Feature vectors were split into primary dataset training, validation and testing splits stratified by family, and the downstream classifier architecture, optimisation settings and per-feature-set hyperparameters are given in Appendix S1 (Figure S2).

For the second criterion, compactness and redundancy, we compared MorphQ, PCA and small-sample DeepClassifier features under the same 512-dimensional output size, asking whether among image morphological variation was concentrated in a smaller subset of informative feature dimensions and whether different feature dimensions carried largely independent information rather than redundant linear signals. We therefore examined three properties of the feature dimensions: their magnitude, their among image variation, summarised as the standard deviation of each dimension across images, and their pairwise Pearson correlation coefficients, for which distributions centred near zero indicate lower linear redundancy. Because the three feature spaces differ in absolute scale, each dimension’s standard deviation was expressed relative to the mean feature value of that method for axis alignment only; the calculation steps are given in Appendix S1 (Figure 2b, e-g).

**Figure 2.**
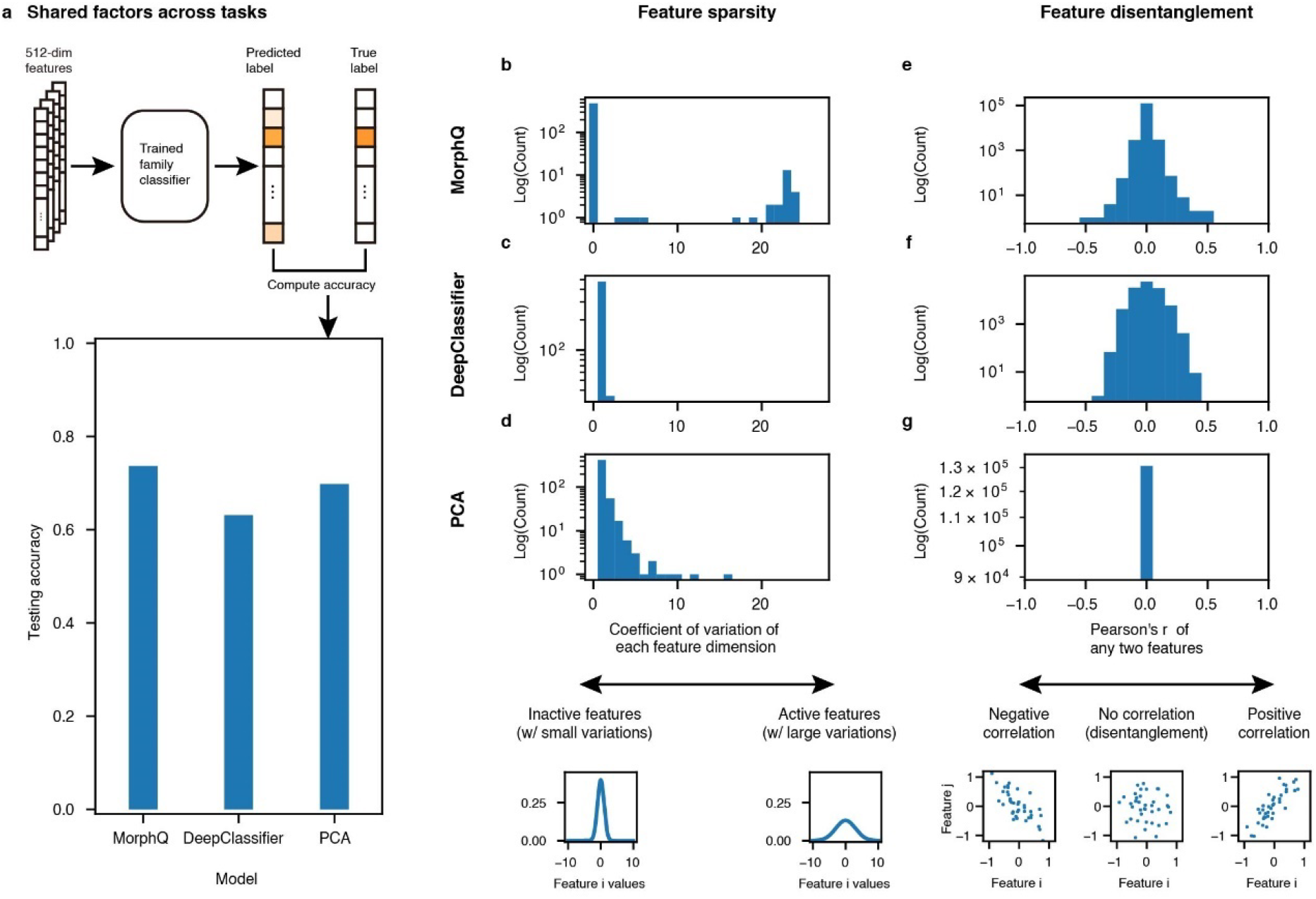
Compactness and downstream utility of MorphQ-derived morphological features. (a) Diagnostic family-classification task; bars show top 1 testing accuracy for classifiers trained on each feature set. (b-d) Frequency distributions of feature variability in MorphQ, DeepClassifier and PCA, with each dimension’s standard deviation scaled per method by the mean feature value for horizontal-axis alignment only. (e-g) Frequency distributions of pairwise correlations among feature dimensions.

For the third criterion, applicability beyond the fitted data and preservation of biologically meaningful similarity, we examined whether family-level structure in the extracted features remained coherent for species not represented in the feature extractors’ fitted or training data. As a prerequisite, we first confirmed that families formed coherent clusters within each feature space, since the subsequent evaluation on isolated-dataset species is only meaningful if family structure is already present. Using the primary dataset, we projected each feature space to two dimensions with Uniform Manifold Approximation and Projection (UMAP) (McInnes et al., 2018), a non-linear ordination method, and quantified family-level clustering from the original 512-dimensional vectors with Silhouette, Calinski-Harabasz, and Davies-Bouldin scores, using family identity as the cluster label. We then measured, for each test image from the isolated dataset, the Pearson correlation between its feature vector and the centroid of its corresponding family calculated from primary-dataset images, analysed these similarities with mixed-effects models including family and image ID as random factors, and added a complementary nearest centroid test reporting top 1 and top 5 accuracies; the post hoc comparison procedure and the dimension-wise inspection of these correlations are given in Appendix S1 (Figure 4).

For the fourth criterion, smoothness and visual interpretability, we evaluated whether the feature space could be rendered as a human inspectable morphospace and whether interpolations across feature space produced gradual morphological transitions rather than abrupt or implausible changes. We used the MorphQ decoder to visualise individual feature dimensions, interpolations across feature space, gradients from downstream analyses and feature combinations that did not necessarily correspond to sampled specimen images. Individual dimensions were interpolated within the range of the empirical feature distribution while the remaining dimensions were held at zero, and four visually distinct specimens were placed at the corners of a 5 × 5 interpolation grid (Figure 3). We also decoded the feature combinations that the downstream family classifier relied on, identified by gradient-based voting across feature dimensions and summarised with PCA into three axes (Figure S3b). PCA served as the decoder-like linear baseline, using the same one axis interpolation and 5 × 5 grid logic (Figure 3c, d); DeepClassifier features could not be evaluated in this way because the architecture did not include an inverse projection mechanism, so Grad-CAM (Selvaraju et al., 2017) is reported only as a local explanation reference and not as a smoothness evaluation (Figure S4). Smoothness was assessed qualitatively because there is no independent quantitative metric for visual similarity among specimens in this context. Full procedures, including the N sweep used to select key feature dimensions, are given in Appendix S1.

**Figure 3.**
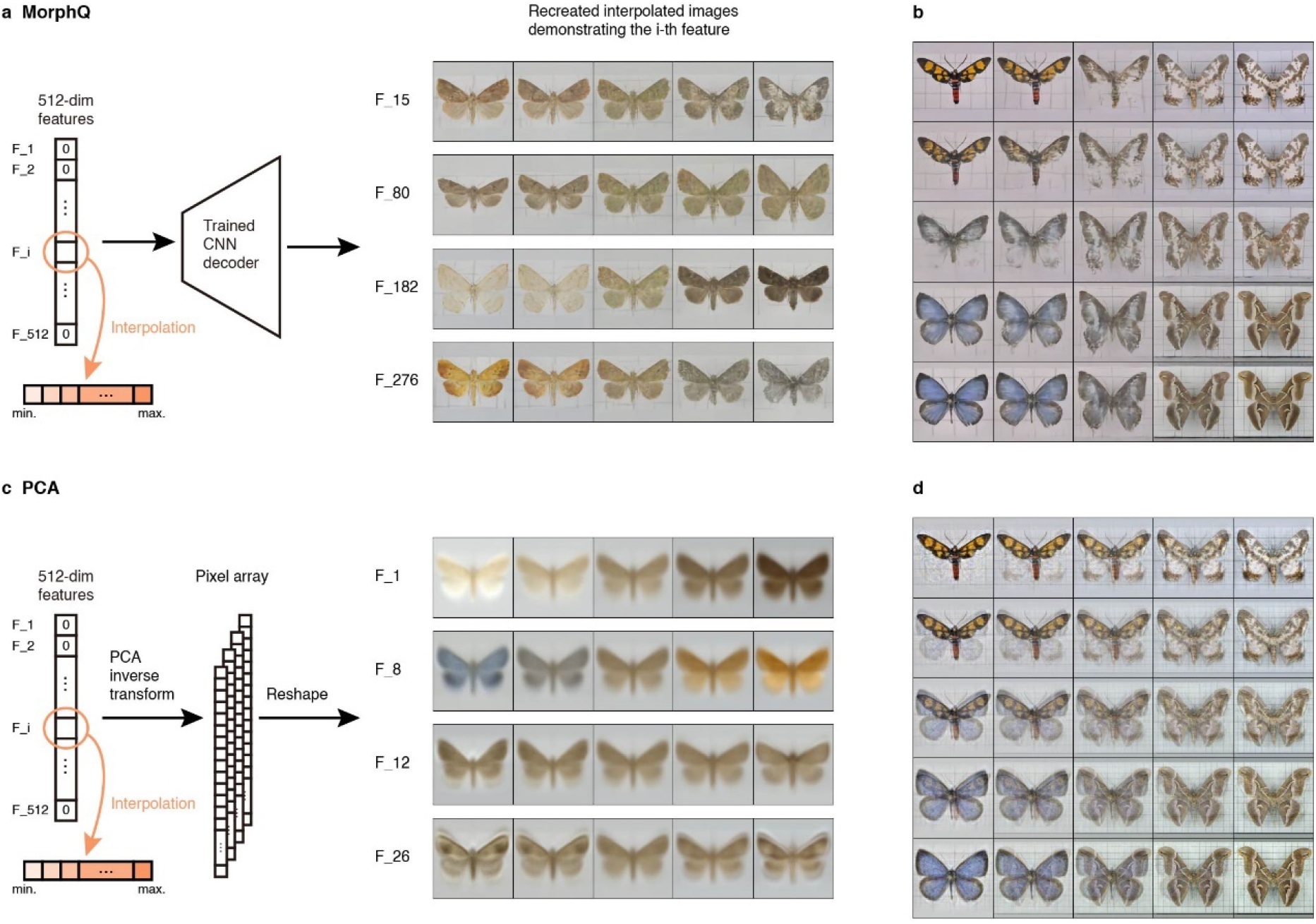
Visual interpretation and generation from MorphQ-derived feature space. (a, c) Workflow and examples for trait-axis visualisation in MorphQ and PCA. (b, d) Interpolated images showing transitions among four real Lepidoptera images in MorphQ and PCA feature spaces; intermediate points are derived feature combinations, not sampled specimens.

### Case Study 1: species-level traits in relation to elevation

We present this analysis as an example, not as a comprehensive explanation, of elevational morphological variation in Lepidoptera. Its purpose was to demonstrate how MorphQ-derived features can be aggregated from images into species-level trait vectors, linked to environmental metadata and decoded into visual morphology for biological interpretation.

We extracted 512-dimensional MorphQ features from all images in the primary dataset and averaged the image-level vectors within species. Species lacking mean elevation data were removed (309 species), leaving 1,559 species. We then applied a separate PCA ordination to the species-level MorphQ vectors and used the first three principal components (PCs) as summary morphological axes for modelling and visualisation. For each of the first three principal components, we fitted a mixed-effects model with the species-level PC score as the response variable, species mean elevation as the fixed effect and family as a random effect. To interpret the PCA-derived morphological axes, we interpolated PC1, PC2 and PC3 within the range of the data distribution, inverse-transformed these positions back into the 512-dimensional MorphQ feature space and decoded them into visual representations with the MorphQ decoder.

### Case Study 2: assemblage-level functional trait diversity

We present the family-level diversity analysis as an example showing how MorphQ-derived morphology can be integrated with trait probability density analyses. Here, “assemblage-level” is used in an operational sense: families were treated as comparable species sets for demonstrating a trait-probability-density workflow, rather than as local ecological communities sampled from shared sites.

We analysed five common Lepidoptera families: Erebidae, Geometridae, Noctuidae, Crambidae and Drepanidae. We extracted 512-dimensional MorphQ features from all images in the primary dataset and applied a separate PCA ordination to reduce these image-level feature vectors to three dimensions, making trait probability density estimation and visualisation tractable. For each species, we calculated the mean and standard deviation of its image-level scores along these three morphological axes. Species-level trait probability densities were estimated with the TPDsMean function in the R package TPD and family-level densities with TPDc (Carmona et al., 2019), with each species assumed to contribute equally to the estimate for its family. We calculated functional richness, functional evenness, functional divergence and relative redundancy (Mason et al., 2005; Carmona et al., 2019), and used rarefaction by sampling different numbers of species 1,000 times for each family (Walker et al., 2008; Ricotta et al., 2012).

Estimated family-level trait probability densities were visualised in the three-dimensional MorphQ-PC space as scatter plots of points sampled from those densities, with contours showing the 95% probability density region. We then used the MorphQ decoder to make these family-level morphospaces visually interpretable, sampling equally spaced points in the PC1–PC2, PC1–PC3 and PC2–PC3 planes, transforming them back into the 512-dimensional MorphQ feature space and decoding them into representative images. These decoded images should be interpreted as visualisations of morphospace positions rather than as sampled specimen images.

## Results

### MorphQ produced a morphospace that can be analysed and visually inspected

After training, MorphQ represented each standardised Lepidoptera image as a 512-dimensional feature vector while retaining a decoder-based route from numerical features back to visual morphology (Figure 1; Figure S1). Consistent with the sparse regularisation objective, many feature dimensions remained close to zero, whereas a relatively small subset varied among specimens.

Each specimen was located as a point in this feature space, and any specified coordinate could be passed to the decoder to generate an image. The decoder reproduced recognisable specimen images from their feature vectors and also generated coherent images for coordinates that did not correspond to any sampled specimen (Figure 1; Figure S1). Such decoded images therefore represent positions in the learned feature space, including unsampled or hypothetical forms, rather than photographs of collected specimens, linking the outputs of downstream analyses back to human-inspectable morphology.

### MorphQ-derived features support downstream reuse under sparse labels

In the downstream family-classification probe, the classifier trained on MorphQ-derived features achieved the highest top 1 accuracy under the two-image-per-species setting of the primary dataset (MorphQ = 0.737, PCA = 0.698, DeepClassifier = 0.631; Figure 2a). Because MorphQ was trained without taxonomic labels, this result indicates that its self-supervised feature space retained family-relevant morphological variation that could be reused by an independent downstream task.

When the same downstream task was instead trained on features from the large-sample DeepClassifier reference, which used 68,913 labelled images, top 1 accuracy rose to 0.791 (Figure S2). Thus, supervised feature learning can perform strongly when extensive labelled data are available. However, MorphQ achieved comparable downstream utility using only the primary dataset of 3,736 images and required no labels. The comparison supports MorphQ as a practical feature extraction framework for ecological and evolutionary analyses where labelled images are limited, unevenly distributed or not aligned with the intended downstream questions.

### MorphQ produced compact and largely non-redundant feature dimensions

All three methods produced 512-dimensional feature vectors. However, their effective use of these dimensions differed. MorphQ showed a clear double-peaked distribution of feature variability (Figure 2b): one group of dimensions remained close to zero across specimens, whereas another varied substantially among specimen images. PCA and DeepClassifier did not show the same separation between near-zero and variation-carrying dimensions (Figure 2c,d). The double-peaked pattern indicates that MorphQ concentrated specimen-level variation into a smaller subset of dimensions rather than distributing it across the full feature space.

Pairwise correlations among feature dimensions also varied by method (Figure 2e-g). PCA had the lowest inter-dimensional correlations, as expected from its orthogonal transformation mechanism. MorphQ showed lower inter-dimensional dependency than DeepClassifier, although not as low as that of PCA. Together, these results indicate that MorphQ produced a feature space with fewer variation-carrying dimensions than the comparator methods while maintaining relatively low redundancy among dimensions, which makes the feature space more tractable for downstream statistical modelling.

### MorphQ converts feature space analyses into visual morphology

When we varied single dimensions across their empirical ranges, the decoded images showed that individual dimensions often captured co-patterned combinations of traits rather than single manually defined descriptors. For example, as the 182nd MorphQ feature dimension varied across its empirical range, the decoded images changed from lighter to darker wing colour, and from a more square outline to a shorter, wider form, accompanied by the disappearance of wing stripes (Figure 3a). A conventional PCA of human-defined measurements can also summarise several traits along one axis, but such a combined axis is hard to turn into a concrete, inspectable image; with the MorphQ decoder this visualisation is produced directly.

MorphQ also produced interpretable transitions and derived forms in feature space. With feature vectors from four distinct specimens at the corners of an interpolation grid, the decoded images changed gradually across the grid, and intermediate positions corresponded to coherent intermediate morphologies rather than to weighted pixel value mixtures (Figure 3b). More generally, the decoder can be applied to any given feature vector or feature value combination, including values derived from sampled specimens, averaged groups, statistical analyses or hypothetical regions of morphospace.

The feature dimensions that most influenced family prediction (the union of each family’s top N = 12 most-voted dimensions) were summarised with PCA into three axes. Decoding the family centroids recovered the representative appearance of each family (Figure S3a), and decoding interpolations along the three PC axes visualised the key morphological variation the classifier relied on (Figure S3b). Across these decoded images, wing shape, colour, lightness and pattern varied in clear and gradual transitions rather than abrupt changes. Grad-CAM showed that the supervised classifier attended mainly to the wings, indicating where the model looks rather than which morphological features drive its prediction (Figure S4).

### MorphQ produced a morphospace with biologically meaningful structure

We evaluated whether MorphQ organised specimen images into a morphospace that reflected biological similarity, using taxonomic family membership as a coarse biological reference. Among primary dataset training images, UMAP visualisations showed clearer family-associated structure for MorphQ and PCA features than for DeepClassifier features (Figure 4a-c). Clustering metrics calculated from the 512-dimensional feature vectors differed in their rankings: DeepClassifier had the highest Silhouette score, whereas MorphQ had the highest Calinski-Harabasz score and the lowest, and therefore best, Davies-Bouldin score (Figure 4d-f). Taken together, they indicate that MorphQ produced a morphospace in which family-associated structure was evident.

**Figure 4.**
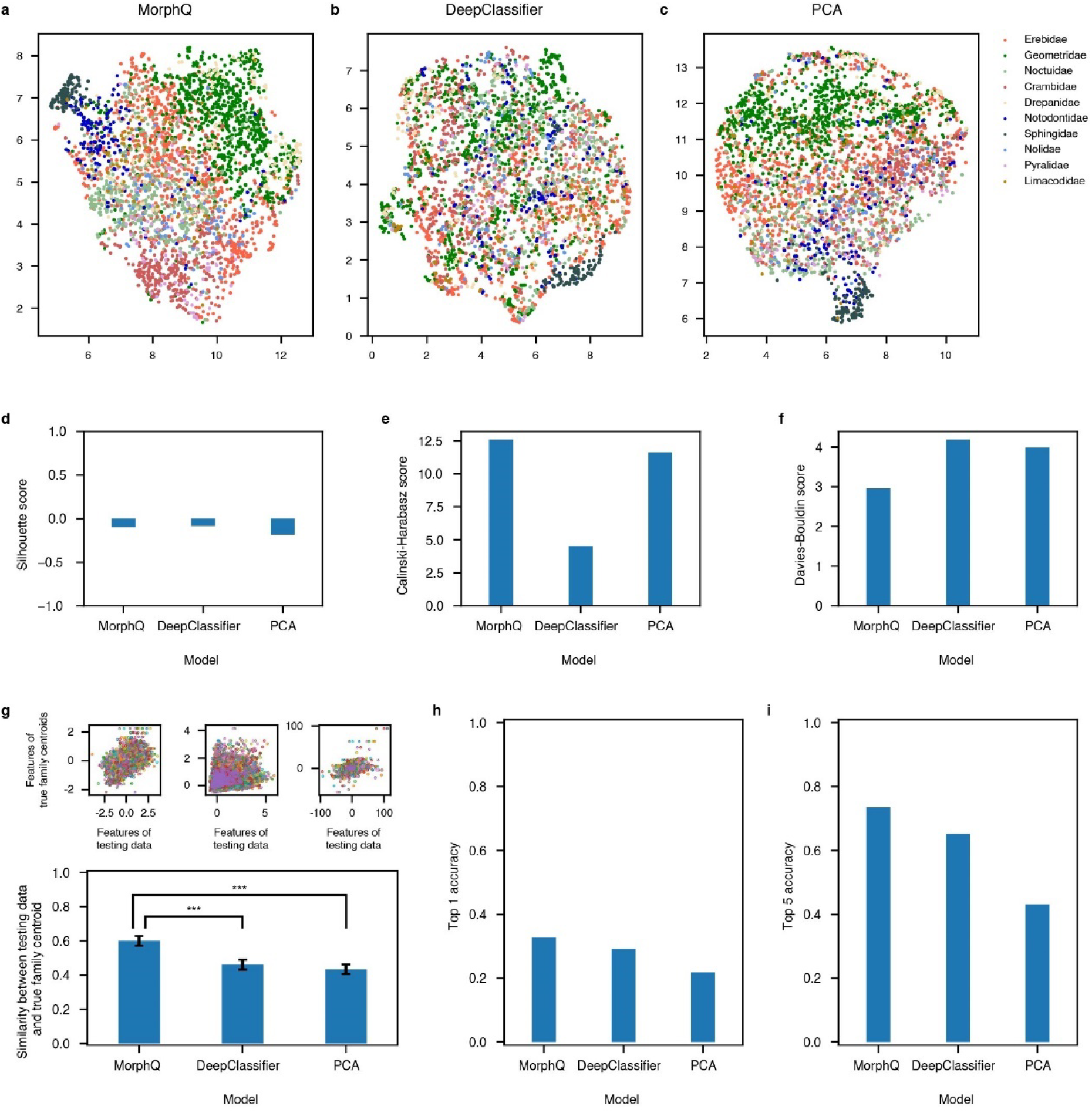
Preservation of biologically meaningful similarity in MorphQ morphospace. (a-c) UMAP visualisations of MorphQ, DeepClassifier and PCA features. (d-f) Clustering metrics using family identity as cluster labels. (g) Similarity between isolated-dataset images and the centroids of their true families. (h, i) Top 1 and top 5 nearest-centroid accuracies for the same test images.

The higher Silhouette score for DeepClassifier should be interpreted cautiously because Silhouette uses sample-to-sample distances and can be more sensitive to extreme or boundary samples, whereas Calinski-Harabasz and Davies-Bouldin scores more directly reflect between-family separation and within-family cohesion. Although the three metrics did not agree on a single ranking, MorphQ performed well across all of them, particularly on the two measures emphasising between-family separation and within-family cohesion.

For isolated-dataset images, MorphQ-derived features showed stronger similarity to the correct family centroid than comparator features did (P < 0.001; Figure 4g; Table S1). Post hoc comparisons confirmed this advantage against both comparators (PCA − MorphQ = −0.17, SE = 0.013, t = −12.62, P < 0.001; DeepClassifier − MorphQ = −0.14, SE = 0.013, t = −10.53, P < 0.001), whereas PCA and DeepClassifier did not differ significantly (DeepClassifier − PCA = 0.03, SE = 0.013, t = 2.09, P = 0.092; Table S1). For MorphQ, isolated-dataset image feature values showed a clear positive association with the corresponding true family centroid values, suggesting that centroid similarity was distributed across feature dimensions rather than dominated by a single feature. The biological structure captured by MorphQ therefore extended beyond the species used during representation learning and remained detectable for isolated-dataset species.

In a nearest-centroid assignment using the same training-derived family centroids, MorphQ-derived features produced the highest top 1 and top 5 accuracies for specimen images from the isolated dataset (Figure 4h,i). Distances in the MorphQ morphospace therefore remained aligned with family structure even for species outside the primary dataset; these accuracies should be interpreted as evidence of distance alignment rather than as a classification result.

### Case Study 1: species-level morphological traits along elevational distributions

Mixed-effects models related the first three species-level MorphQ axes to mean elevation, with family as a random effect (Figure 5a). PC1 was not significantly related to species mean elevation (Figure 5b). In contrast, PC2 showed a significant negative relationship with elevation (Figure 5c) and PC3 a significant positive relationship (Figure 5d), and both held after controlling for family-level non-independence. Part of the species-level MorphQ morphospace therefore covaried with elevation, whereas its other major axes did not.

**Figure 5.**
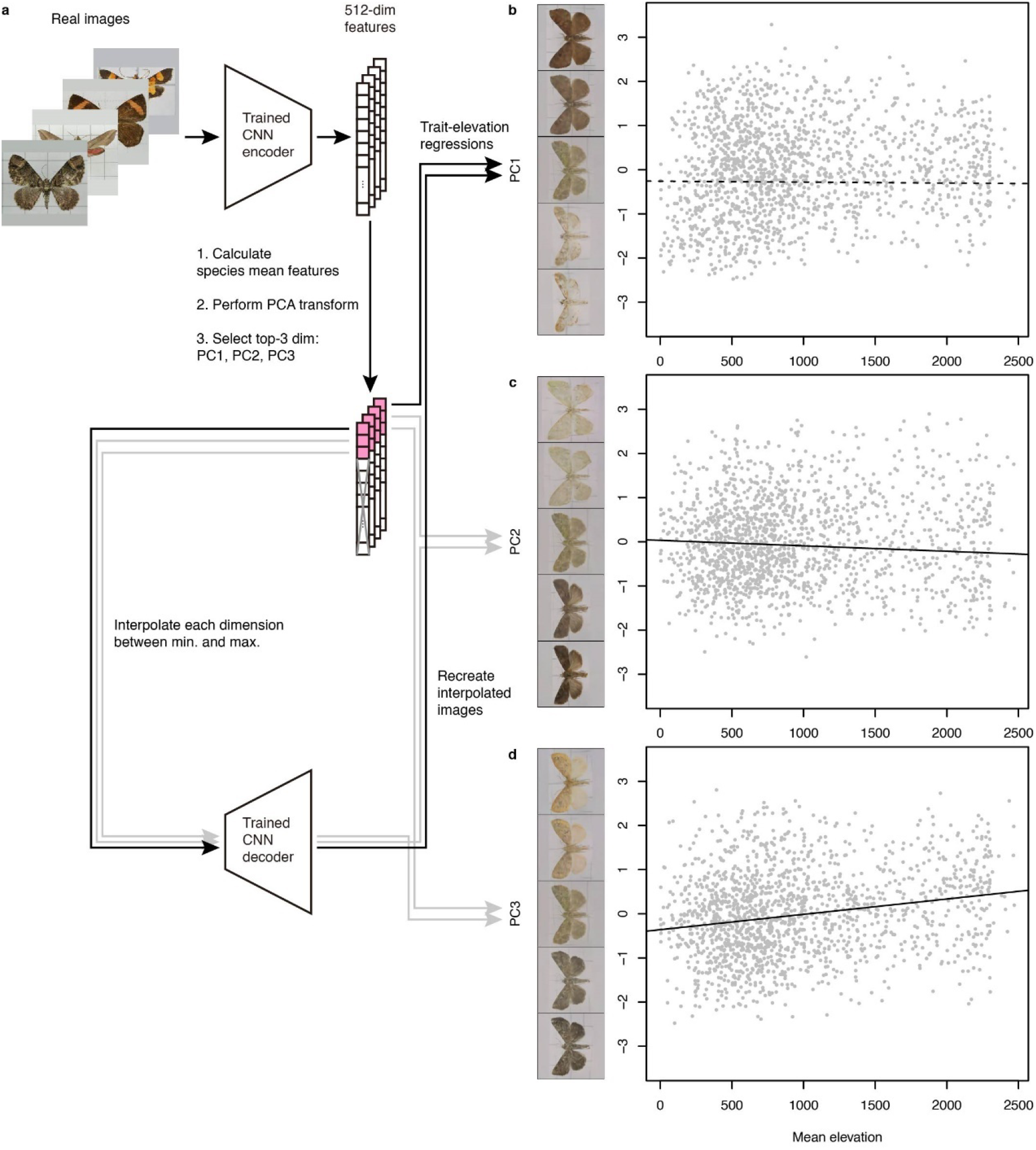
Case study of species-level trait-environment analysis using MorphQ. (a) Workflow for the species-level trait–environment analysis. (b-d) Species-level morphological axes after dimensionality reduction in relation to elevational distribution, with decoder-generated images of the morphology on each axis.

Decoding the associated axes made these relationships interpretable as morphology. Along the portions of PC2 and PC3 associated with higher mean elevation, the decoded images showed stronger contrast between forewings and hindwings, with forewings darker than hindwings (Figure 5c,d). MorphQ renders the elevation-associated axes as a specific, inspectable wing contrast gradient rather than an abstract statistical direction.

### Case Study 2: assemblage-level functional diversity in morphospace

We compared functional diversity among five common Lepidoptera families in MorphQ morphospace (Figure 6a). After rarefaction, Erebidae still had significantly higher functional richness than Geometridae despite their similar species numbers in the dataset (Figure 6b). Functional evenness did not differ significantly among families (Figure 6c). Drepanidae showed the highest functional divergence, indicating a higher proportion of species occupying distinctive regions of morphospace, whereas Crambidae had the highest relative redundancy and Drepanidae the lowest (Figure 6d,e).

**Figure 6.**
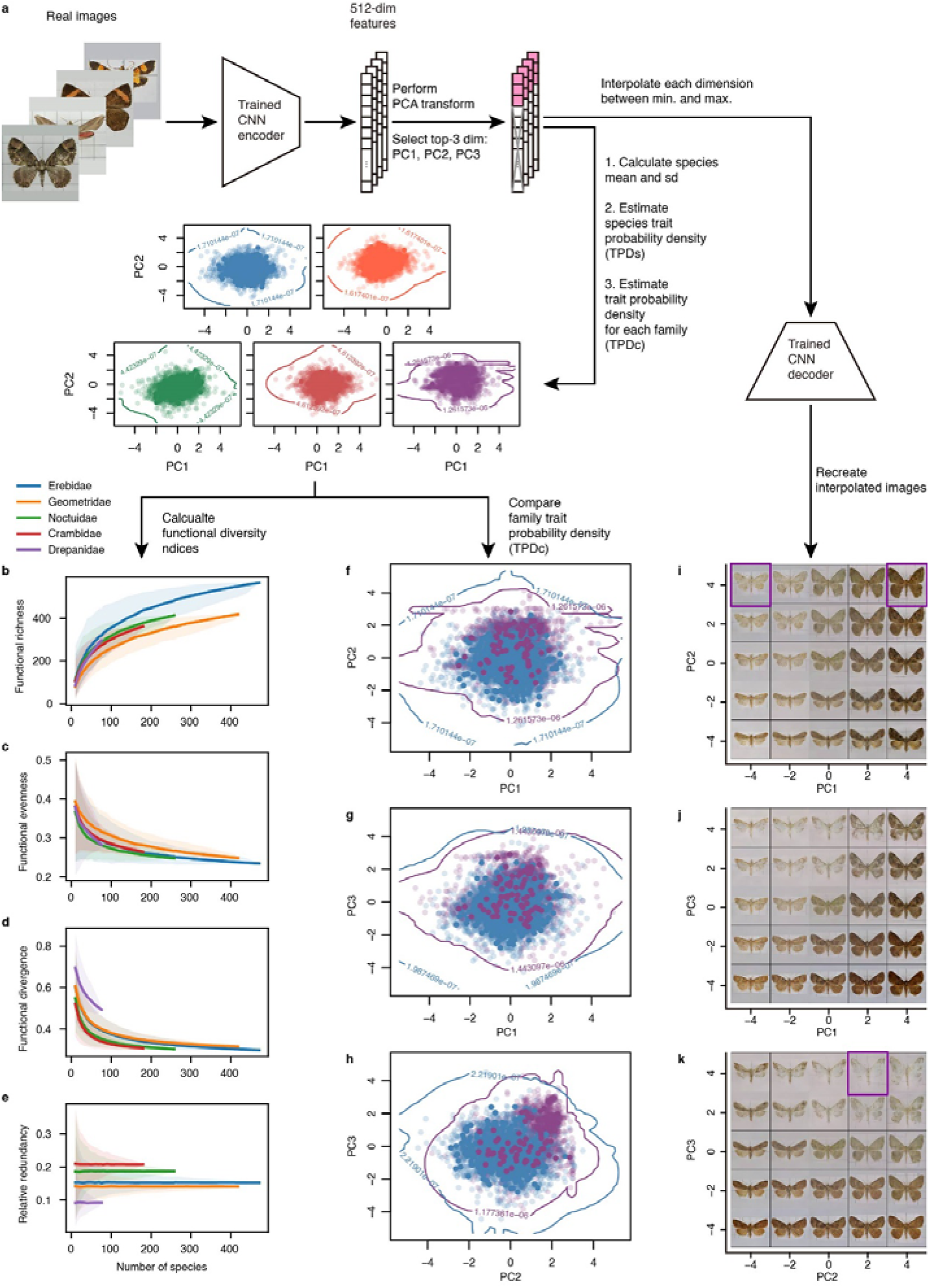
Case study of assemblage-level functional diversity using MorphQ. (a) Workflow for the family-level functional diversity analysis. (b-e) Functional richness, evenness, divergence and relative redundancy for five common Lepidoptera families. (f-h) Trait probability density distributions in MorphQ morphospace. (i-k) Decoder-generated images of morphology at sampled positions in morphospace, including derived positions not corresponding to sampled specimens.

Compared with Erebidae, Drepanidae showed more localised high density regions in MorphQ morphospace, consistent with its high functional divergence (Figure 6f-h). Decoder visualisations of the PC1-PC2, PC1-PC3 and PC2-PC3 planes then showed which morphological forms occupied different regions of the morphospace (Figure 6i-k). Comparing family distributions with the decoded morphology distinguished regions of shared morphology, visually distinctive forms and combinations implied by the morphospace but not represented among the sampled specimens.

## Discussion

### MorphQ provides a reusable and visually inspectable morphospace framework

MorphQ provides a self-supervised framework for transforming organismal images into quantitative and visualisable morphological features. Its central contribution is the integration of three components that are often separated: label-free feature learning, a compact feature space for statistical analysis, and a decoder that can translate feature-space positions back into visual morphology. It addresses a key challenge in morphology-based ecology and evolution: how to derive feature-rich and reproducible representations of complex form without first deciding which individual colours, shapes, or patterns must be measured.

Our empirical results support this framing: MorphQ-derived features were compact, relatively non-redundant, reusable in a downstream diagnostic task, and retained biologically meaningful similarity for species absent from model training. The reconstruction objective encourages the model to retain image-wide visual structure rather than only label-relevant features, while sparse regularisation favours a feature space in which many dimensions remain near zero. The decoder then makes this feature space inspectable, generating images from individual dimensions, feature combinations, group centroids, interpolation paths and statistical gradients so that statistical results can be examined as morphology.

MorphQ therefore differs from both manual trait extraction and standard supervised image classification. Manual approaches are valuable when researchers have strong prior hypotheses, but they necessarily represent only the descriptors selected in advance. Supervised classifiers can be highly effective for prediction tasks, especially when many labelled images are available, but their learned features are optimised for a specific prediction target and may be difficult to reuse or interpret as general morphology. MorphQ instead learns a morphospace without taxonomic labels or predefined descriptors, and then allows researchers to ask downstream ecological or evolutionary questions within that space.

### Scope, assumptions and recommended practice

The analyses here used standardised Lepidoptera specimen images, cropped, scaled and padded into a consistent image format. This setting tests whether MorphQ can learn and visualise morphological features from comparable specimen images, but does not establish that the same model or workflow will perform equally well on non-standardised images such as citizen-science photographs. Applying MorphQ to such images would require additional preprocessing or domain-specific training data to ensure the feature space represents morphology rather than image context.

MorphQ features should therefore be treated as image-derived trait candidates until dataset-specific checks support their biological interpretation. Users should examine image standardisation, segmentation and reconstruction quality, taxonomic and sampling balance, and effects of background, pose, lighting and image source, and should test whether distances, clusters and gradients are stable across training runs and preprocessing choices. These checks are especially important before interpreting feature axes as ecological or evolutionary trait gradients, because apparent structure may reflect image acquisition or sampling rather than morphology.

The broader framework is not restricted to the particular implementation used here. Recent fine-grained image classification methods (Akata et al., 2015; Xiao et al., 2015), non-autoencoder self-supervised methods (Jaiswal et al., 2021), metric-learning approaches (Schroff et al., 2015; Chen et al., 2020), and generative classifiers (Grathwohl et al., 2020; Yang et al., 2021) provide possible routes for extending the same conceptual framework to less standardised images. For instance, Hoyal Cuthill et al. (2019) combined triplet-loss deep classifiers with butterfly phenotype analysis to extract comparable image-derived features, illustrating how such approaches can be applied to organismal morphology. However, such extensions must preserve two requirements: distances and directions in feature space must remain biologically comparable, and analysed positions in feature space must remain visually interpretable. If feature vectors are dominated by background, pose, or illumination, the resulting morphospace will not support meaningful ecological or evolutionary inference. If the visualisation component is absent or unreliable, the framework may still provide image embeddings, but it will lose the decoder-based link that allows statistical results to be inspected as morphology.

### Methodological implications for ecology and evolution

Deep learning has already reduced labour in species identification, object detection and animal pose tracking (Norouzzadeh et al., 2018; Høye et al., 2021; Graving et al., 2019; Lauer et al., 2022). MorphQ applies representation learning to a different problem: deriving quantitative representations of morphology that can be analysed statistically and inspected visually. MorphQ thereby contributes to the long-standing effort to construct morphospaces for complex organismal forms, with less dependence on predefined trait choices while keeping axes, gradients and group summaries linked to inspectable images.

Our two case studies illustrate this methodological role at two different analytical scales. In the elevation analysis, MorphQ linked species-level morphological axes to environmental metadata and translated associated axes into visual gradients of wing morphology. In the family-level functional diversity analysis, MorphQ connected functional richness, divergence, and redundancy metrics to regions of morphospace and to decoded morphologies that made those regions visually interpretable.

Image-derived features can be useful for modelling, but without a route back to morphology their biological interpretation is hard to evaluate. The decoder-based visualisation supplies that route: it does not replace biological validation, but it lets researchers see the morphological patterns implied by statistical axes, clusters or gradients, which distinguishes MorphQ-derived trait analyses from analyses based only on opaque embeddings. Decoder-generated images can also serve as interpretable intermediate outputs for targeted follow-up measurements of shape, colour, brightness or pattern, connecting MorphQ axes back to predefined descriptors. MorphQ therefore provides a reproducible framework for constructing visualisable morphospaces from standardised specimen images, making complex image-derived morphology available for downstream ecological and evolutionary analyses when predefined descriptors are incomplete and labelled data are limited, and yielding visual hypotheses that can be inspected before biological interpretation.

## Data availability statement

An anonymised repository containing the source code, trained model weights, configuration files, example data and scripts required to reproduce the analyses is available at https://anonymous.4open.science/r/MorphQ-ECD4/. Specimen metadata were obtained from the GBIF dataset “Data-set of Moth Specimen from TESRI” (https://doi.org/10.15468/kjjlnf), and the large specimen image dataset was assembled from the Taiwan Moth Information Center (https://twmoth.tesri.gov.tw) as described in Materials and Methods.

## Use of AI tools

Generative AI tools (ChatGPT and Claude) were used for language editing of the manuscript and to assist with drafting and debugging analysis code. AI-assisted code is identified as such in the repository. No AI tool was used to generate data, results or interpretations. The authors reviewed and verified all outputs and take full responsibility for the content of this article.

## Supporting information

Support Information

