## Supplementary material for "MorphQ: label-free quantification and visualisation of complex morphology from standardised specimen images": Support Information

### Supporting information

### Appendix S1. Detailed materials and methods

#### Image datasets and preprocessing — detailed procedures

Non-specimen images and low quality specimen images, such as those with badly broken or missing wings or wings not fully spread, were removed, yielding 68,913 Lepidoptera specimen images.

For this analysis, we treated family as the natural group membership of each isolated-dataset species and compared the feature similarity between each isolated-dataset species and the centroid of its corresponding family calculated from the primary dataset.

In our implementation, we trained a YOLOv4 model (Bochkovskiy et al. 2020) to predict specimen bounding boxes and crop the target specimen from each image. This cropping step was used only to standardise image framing and is not specific to YOLOv4; other contemporary pretrained object detection or semantic segmentation models could be substituted for this step without task-specific model fine tuning, provided that they can identify the focal specimen consistently. The width and height of each bounding box were multiplied by 1.1 to leave space between the target specimen and the image border, thereby reducing the risk that wingtips or specimen edges were cropped out. The long side of each cropped image was rescaled to 256 pixels, and the short side was padded to 256 pixels, yielding a 256 × 256 × 3 RGB image. The padding colour was determined separately for each cropped image. Before padding, three 10 × 10 pixel patches were sampled from the top centre, bottom left and bottom right of each cropped image, in regions chosen to avoid the specimen. We then applied K-Means clustering with k = 2 to the sampled RGB values of these patches and used the average RGB value of the larger cluster as the padding value.

**Comparator methods — implementation and training details**

PCA features were obtained by flattening each standardised 256 × 256 × 3 RGB image into a 196,608-dimensional pixel-channel vector and retaining 512 principal components, matching the feature dimensionality used by MorphQ.

The fitted PCA parameters, including the feature means and component loadings, were retained so that 512-dimensional PCA coordinates could be inverse-transformed and reshaped back into 256 × 256 × 3 RGB images.

The pooled representations formed 512-dimensional feature vectors, which were then passed to the classifier head for species prediction during training. After training, these pooled 512-dimensional vectors were used as DeepClassifier-derived features for downstream comparisons. The small-sample DeepClassifier was trained from scratch on the primary dataset using the same image augmentations as MorphQ. The primary dataset was split into training, validation and testing splits at a ratio of 6:2:2, with images of the same species distributed as evenly as possible across splits. Because only two images were available for each species, some species appeared only in the primary dataset training split, whereas others appeared once in either the primary dataset validation or testing split. The small-sample DeepClassifier training used a batch size of 256 and Adam optimisation with a learning rate of 5 × 10−5 and weight decay of 1 × 10−5. Early stopping was based on validation loss, with patience set to 100 epochs. The model with the minimum validation loss was used for downstream analyses. This training run stopped at 506 epochs.

The large-sample DeepClassifier reference used the same architecture and training procedure as the small-sample DeepClassifier but was trained with the large-sample reference dataset of 68,913 labelled images and different Adam hyperparameters: learning rate 8 × 10−5 and weight decay 2 × 10−5. This training run stopped at 202 epochs.

**Evaluation of image-derived morphological representations — detailed procedures**

For the downstream family classifier probe, feature vectors were split into primary dataset training, validation and testing splits stratified by family, at an approximately 6:2:2 ratio as evenly as possible. The classifier consisted of two hidden layers and a 47-class output layer corresponding to the families in the dataset. Each hidden layer included a 50% dropout layer, a fully connected layer with 256 neurons and a leaky ReLU activation with negative slope = 0.2. The output layer was a fully connected layer with 47 neurons, followed by a leaky ReLU activation; the final softmax step was handled by PyTorch cross-entropy loss. Weights of the downstream family classifiers were initialised with PyTorch defaults. For each extracted feature set, including MorphQ-derived, PCA-derived, small-sample DeepClassifier-derived and large-sample DeepClassifier-derived features, a family classifier with the same architecture was trained independently on the primary dataset feature split using Adam optimisation with batch size set to 3,000. Because the training split contained 2,390 primary dataset images, all training samples were processed in a single batch within each epoch. Hyperparameters were selected separately for the family classifier trained on each feature set to keep validation accuracy close to training accuracy while minimising validation loss: learning rate = 1 × 10−4 and weight decay = 8 × 10−6 for MorphQ-derived features; learning rate = 5 × 10−4 and weight decay = 5 × 10−5 for PCA-derived features; and learning rate = 1 × 10−3 and weight decay = 5 × 10−5 for DeepClassifier-derived features. For each feature set, the final downstream family classifier model was selected as the checkpoint with the minimum validation loss, which occurred at epoch 1,616 for MorphQ-derived features, epoch 79 for PCA-derived features, epoch 69 for small-sample DeepClassifier-derived features, and epoch 158 for large-sample DeepClassifier-derived features.

For each method, we summarised the among image variation of every feature dimension as the standard deviation of that dimension across images. Because the three feature spaces differ in absolute scale, we expressed each dimension's standard deviation relative to the mean feature value of that method (a per method coefficient-of-variation scaling) solely so that the three distributions could be placed on a common horizontal axis for visual comparison; this scaling affects axis alignment only and not the shape of the distributions on which our interpretation rests. For MorphQ, whose latent prior is centred on zero, dimensions that remained close to zero across images and showed little variation were interpreted as contributing little to among image morphological variation, whereas dimensions that departed from zero and varied among images were interpreted as contributing to among-image morphological variation. To evaluate redundancy, we calculated Pearson correlation coefficients between all pairs of feature dimensions. Correlation distributions centred near zero indicated lower linear redundancy among feature dimensions.

For each method, family centroids were calculated by averaging feature vectors from primary-dataset images within each family. For each isolated-dataset image, we measured the Pearson correlation between its feature vector and the centroid of its corresponding family. These similarities were analysed using mixed-effects models, with family and image ID (the filename-based unique image identifier) included as random factors, followed by post hoc comparisons. To examine whether this centroid similarity was distributed across feature dimensions rather than driven by a few dimensions, we inspected, for each isolated-dataset image, the dimension-wise relationship underlying this Pearson correlation by plotting its 512 per-dimension feature values against the corresponding true-family centroid values. As a complementary nearest centroid test, we assigned each isolated-dataset image to the closest family centroids in Euclidean distance and calculated top 1 and top 5 nearest centroid accuracies.
To visualise individual MorphQ feature dimensions and their smoothness, each dimension was interpolated within the range of the empirical feature distribution while the other dimensions were set to zero, the centre of the latent prior, and the resulting 512-dimensional vectors were decoded into images. To further evaluate smoothness in MorphQ, we selected four visually distinct Lepidoptera images based on wing shape, wing colour, stripe pattern and taxonomic identity. Their MorphQ feature vectors were placed at the four corners of an interpolation space, and intermediate 512-dimensional feature vectors were obtained by interpolation. These vectors were then decoded to produce a $5 \times 5$ MorphQ image grid, in which the corner images

corresponded to the four selected specimens and the internal images visualised interpolated positions between them.

Beyond individual dimensions and pairwise interpolations, we also used the decoder to visualise the feature combinations that the downstream family classifier relied on. Key feature dimensions were identified from the downstream family classifier trained on MorphQ features. For each primary dataset image, we backpropagated the predicted class score to the MorphQ feature layer and ranked all 512 feature dimensions by their gradient magnitude. For a given number N, each image contributed a vote to each of its own top N gradient-ranked dimensions, and these votes were tallied separately within each family; for every family we then selected its N most-voted dimensions, and the key feature set for that value of N was the union of these per family selections across all families. We swept N from 1 to 50 and, treating family identity as the cluster label, evaluated how well the corresponding union of dimensions separated families using a combined criterion of Silhouette, Calinski-Harabasz, and Davies-Bouldin scores computed on those original feature dimensions; this procedure selected N = 12. The resulting union of key dimensions was then summarised with PCA into three axes, capturing the directions of greatest variation among the classification-relevant dimensions so that this discriminative structure could be rendered as morphology; the PCA step for the MorphQ key feature visualisation is intended to demonstrate the most prominent directions of morphological variation among the key features rather than to prescribe a required analysis step. Key feature combinations were visualised by interpolating each PCA axis across its empirical value range, inverse-transforming the interpolated coordinates back into the 512-dimensional MorphQ feature space and decoding them into images (Figure S3b).

For the PCA method, its inverse transformation can serve as the decoder-like linear baseline for this criterion and can project retained component values back into image space. We therefore visualised individual PC axes using the same one axis interpolation logic applied to MorphQ feature dimensions, and further tested smoothness using 5 × 5 interpolation image grids. For comparison with the MorphQ image grid, equivalent PCA interpolation image grids were constructed by interpolating PCA feature vectors between the same selected specimens, inverse-transforming the interpolated vectors and reshaping them back into 256 × 256 × 3 RGB images. Smoothness was assessed qualitatively because there is no independent quantitative metric for visual similarity among specimens in this context. DeepClassifier features could not be evaluated for smoothness or feature space visualisation using the same feature-to-image procedure because the DeepClassifier architecture did not include an inverse projection mechanism. As a separate local explanation reference, we used Grad-CAM to identify image regions that influenced the supervised classifier (Selvaraju et al., 2017); this analysis was not treated as a smoothness evaluation (Figure S4).

**Software availability and reproducible workflow**

All Python code was implemented and executed in Python 3.7.13 on Ubuntu 18.04 using two NVIDIA GeForce GTX 1080 Ti GPUs for model training, feature quantification, and visualisation. The main Python libraries were PyTorch 1.8.0 with CUDA 11.1, NumPy 1.19.5, Pandas 1.0.5, SciPy 1.4.1, Scikit-learn 0.22.2.post1, Scikit-image 0.16.2, Matplotlib 3.2.2, ImgAug 0.4.0, and UMAP 0.4.6. Mixed-effects models and trait probability density analyses were implemented in R 4.1.2 using data.table 1.14.2, lme4 1.1.28, MuMIn 1.46.0, lmerTest

3.1.3, emmeans 1.7.2, and TPD 1.1.0. The source code, trained model weights, example data, and a reproducible vignette walking through the full pipeline are available at https://anonymous.4open.science/r/MorphQ-ECD4/.

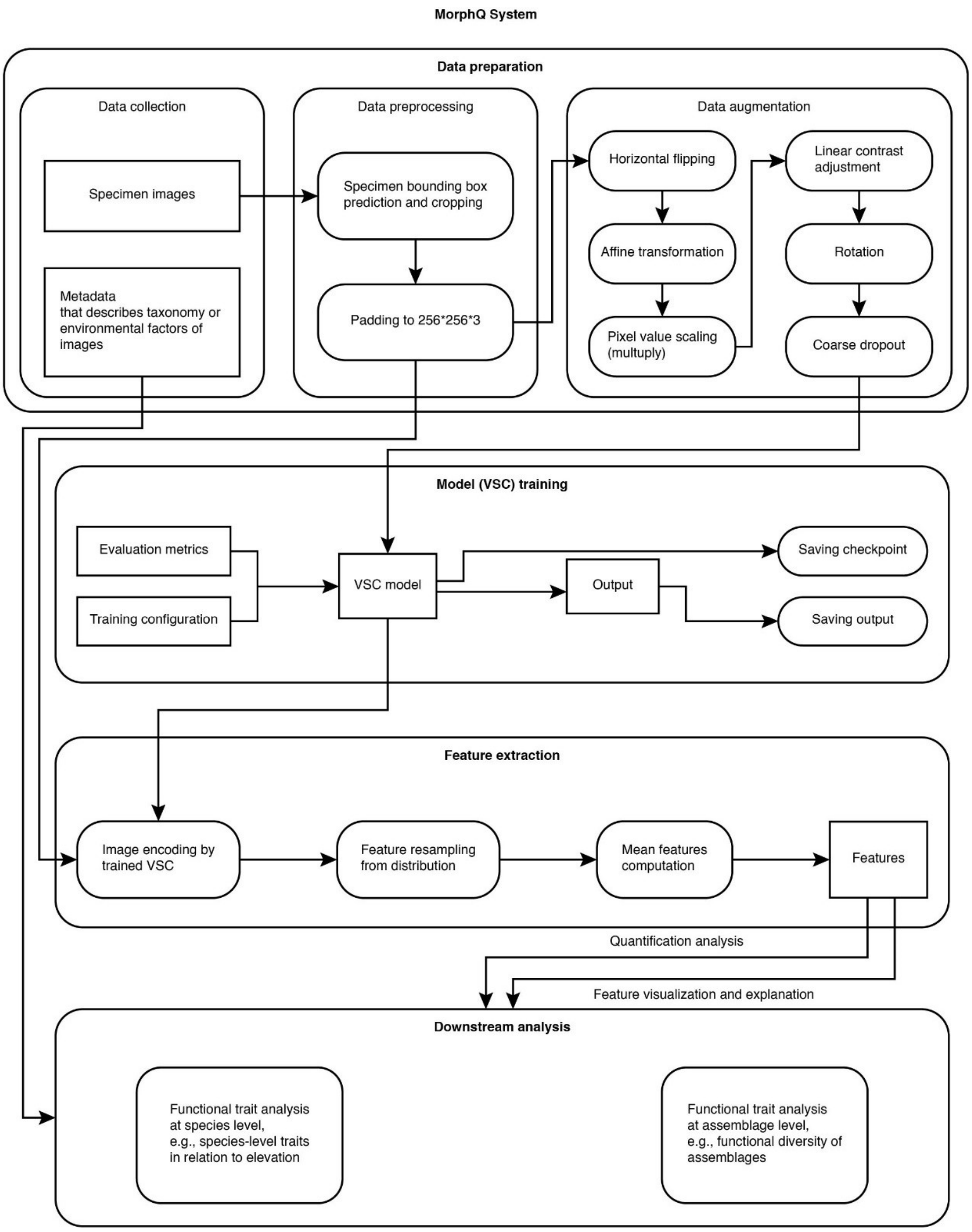


**Figure S1.** Detailed MorphQ workflow for morphological feature quantification and visualisation.

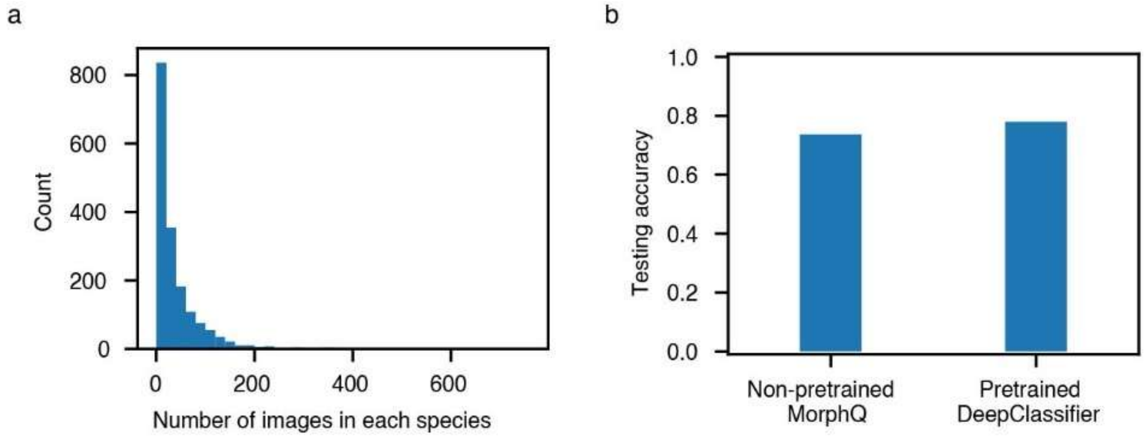


**Figure S2.** Large-sample supervised DeepClassifier pretraining. (a) Frequency distribution of species labels in the large TESRI moth image dataset. (b) Comparison of testing accuracy for downstream family classifiers trained on MorphQ features and on features extracted from the large data pretrained DeepClassifier.

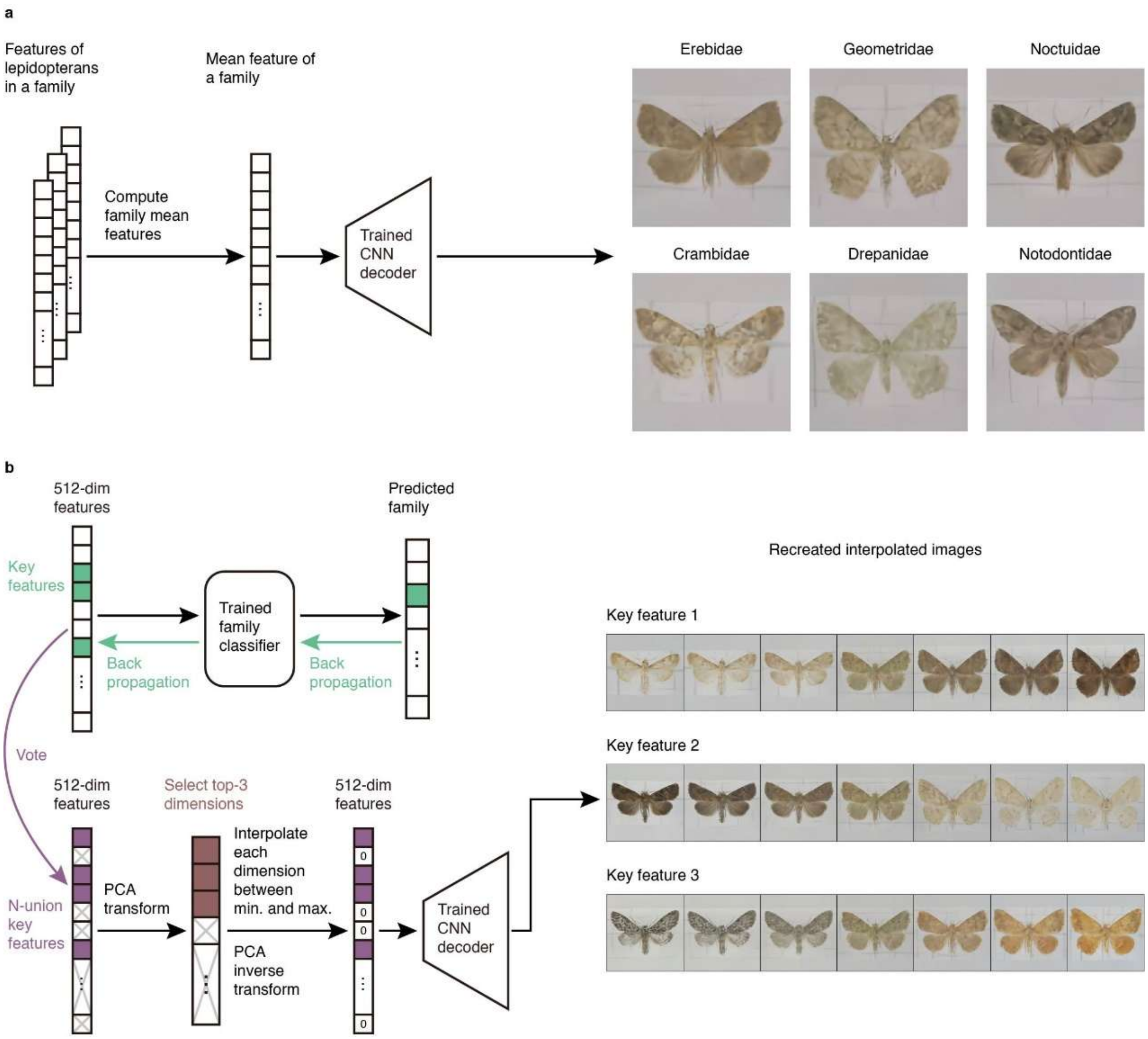


**Figure S3.** Additional decoder-based trait visualisations. (a) Visualisation of family centroids for six common Lepidoptera families. (b) Visualisation of key feature combinations used in the diagnostic family-classification analysis.

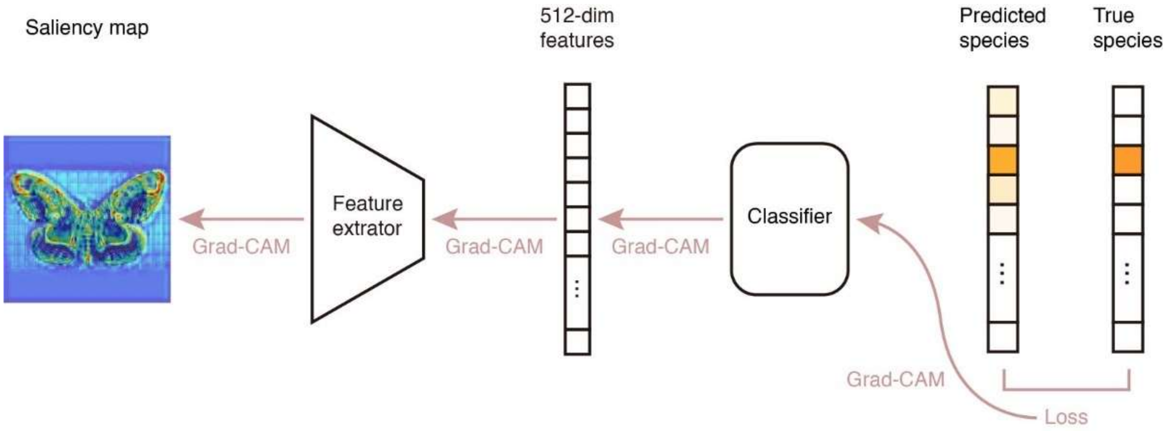


**Figure S4.** Grad-CAM local explanations for the supervised DeepClassifier model.

**Table S1.** Post-hoc comparisons from the mixed model testing the influence of feature extraction method on similarity between test images from the isolated dataset and true family centroids.

| Contrast | Estimate | SE | t.ratio | P |
|---|---|---|---|---|
| DeepClassifier - PCA | 0.03 | 0.013 | 2.09 | 0.092 |
| DeepClassifier - MorphQ | -0.14 | 0.013 | -10.53 | <0.001 |
| PCA - MorphQ | -0.17 | 0.013 | -12.62 | <0.001 |

**Table S2.** Model architecture and hyperparameters.

| **Component** | **Setting** |
|---|---|
| Encoder — input | Instance normalisation |
| Encoder — first convolution | Kernel size = 5, padding = 2, stride = 1; batch normalisation; leaky ReLU with negative slope = 0.2; 32 feature maps at input resolution |
| Encoder — down-sampling | Average pooling with kernel size = 2 |
| Encoder — residual stack | Average pooling down-sampled the representation a further six times; output channels expanded as [64, 128, 256, 512, 512, 512] |
| Residual block | Two convolution layers with kernel size = 3, padding = 1 and stride = 1, with batch normalisation and leaky ReLU; identity connection expanded by a convolution layer with kernel size = 1, stride = 1 and no padding |
| Latent representation | 512-dimensional spike-and-slab distribution (mean, log variance, log spike) |
| Decoder | Mostly symmetric to the encoder; down-sampling layers replaced by up-sampling layers; order of residual block output channels reversed; reconstructed without an instance normalisation layer |
| Weight initialisation | PyTorch defaults |
| Augmentation — flipping | Horizontal flipping with probability = 0.5 |

| | |
|---|---|
| Augmentation — affine scaling | 0.9 to 1.1, applied independently along the horizontal and vertical axes, with probability = 0.9 |
| Augmentation — brightness and contrast | RGB-channel-independent scaling from 0.9 to 1.1 with probability = 0.3 |
| Augmentation — translation | Up to 10% along the horizontal and vertical axes with probability = 0.3 |
| Noise — rotation | Random rotation from −180° to 180° |
| Noise — dropout | Coarse dropout of varying sizes with probability = 1/3 |
| Optimiser | Adam with a learning rate of 0.0002 |
| Batch size | 64 |
| Epochs | 70,850; no formal early stopping rule or predefined epoch limit |
| Framework | PyTorch |
